# Quantitative Detection of Amyloid Fibrils using Fluorescence Resonance Energy Transfer (FRET) Between Engineered Yellow and Cyan Proteins

**DOI:** 10.1101/2024.12.14.628482

**Authors:** Caitlyn Moustouka, George I. Makhatadze

**Author notes:** Corresponding Author: George Makhatadze, Center for Biotechnology and Interdisciplinary Studies, Rensselaer Polytechnic Institute, Troy, NY 12180, USA.

## Abstract

Over 20 human diseases are caused by or associated with amyloid formation. Developing diagnostic tools to understand the process of amyloid fibril formation is essential for creating therapeutic agents to combat these widespread and growing health problems. Here, we capitalize on our recent striking discovery that green fluorescent protein (GFP), one of the most used proteins in molecular and cell biology, has a high intrinsic binding affinity to various structural intermediates along the fibrillation pathway, independent of amyloid sequence. Using engineered GFP with the fluorescence properties of Aquamarine and mCitrine, we developed a FRET-based sensor to quantitatively monitor amyloid fibrils. The proof-of-principle characterization was performed on a test system consisting of PAPf39 fibrils.

## 1. Introduction

Amyloid fibrils are a class of highly ordered protein aggregates with a wide variety of implications across all domains of life. Perhaps most notable is the role of amyloids in numerous human diseases, including Alzheimer’s disease, Parkinson’s disease, Huntington’s disease, and dozens of other amyloidoses, cancers, and dementias (9); cationic fibrils have also been shown to enhance the infectivity of enveloped viruses (56; 33; 18; 6). In addition to their pathological roles, amyloids are involved in diverse functional roles, including biofilm formation and surface adhesion in bacteria (12), prion-based phenotype propagation in fungi (28), and modulation of long-term memory in higher eukaryotes (51). Despite decades of progress, many aspects of amyloidogenesis and the roles of amyloids in disease states are not understood well enough to reliably treat or prevent amyloid diseases.

Identification of amyloid structures is commonly achieved using fibril-specific dyes, such as thioflavin-T (ThT) or Congo red (CR), which undergo a significant change in their fluorescence properties upon binding amyloid, independent of the constituent peptide sequence (2), making them useful in studying a broad range of amyloid systems. Despite their widespread acceptance as key amyloid research tools, an accumulation of recent evidence suggests a lack of ThT and CR amyloid-specificity. Fibril dyes have been shown to recognize amorphous and non-amyloid fibrillar protein assemblies (7; 23; 58), certain native monomeric proteins (24; 45; 1; 10), DNA (32; 40), lipids (29; 13), and other cellular components (31; 39). Higher specificity can be achieved using sequence-specific antibodies or fluorescently labeled amyloid precursors (43; 2), but these methods are inherently non-generalizable across systems with different constituent peptide sequences.

In addition to the lack of specificity displayed by amyloid dyes, they are only able to recognize amyloid structures that exhibit the characteristic cross-β architecture, which excludes prefibrillar oligomers (8; 42). Current evidence suggests that these soluble oligomeric states of amyloids exhibit the most drastic cytotoxic effects through cell membrane permeabilization (15), driving synaptic dysfunction and subsequent neuronal death in amyloid-related neurodegeneration (21). These important early stages of amyloidogenesis are effectively invisible in the context of amyloid dye detection, requiring the use of alternative tools to study these prefibrillar species (20; 19). The generalizability and ease of using amyloid dyes make them favorable research tools, but their utility is constrained by the previously mentioned issues.

We have previously reported that fluorescent proteins (FPs) bind the fibril cores of various disease-related amyloids, including those formed from amyloid-β residues 1-42 (Aβ_42_) and 1-40 (Aβ_40_), α-synuclein, tau, insulin, semenogelin-1 residues 86-107 (SEM1_86-107_), and prostatic acid phosphatase residues 248-286 (PAPf39, otherwise known as “semen-derived enhancer of viral infection,” or SEVI); this interaction is fibril-specific, as FPs show no affinity for amorphous aggregates or non-amyloid fibril structures (58). Additionally, time course confocal microscopy of PAPf39 and SEM1_86-107_ throughout their aggregation suggests FPs recognize transient species early in the aggregation pathway that are not recognized by CR, perhaps being prefibrillar oligomers (58). The presence of FPs within amyloidogenic protein fusion constructs or as individual molecules has been shown to decrease the rates of aggregation of various amyloid systems (53; 3; 5; 58), further supporting an interaction with both early amyloidogenic aggregates and mature fibrils. Taken together, FPs exhibit fibril structure specificity, generalizability across amyloid systems, and recognition of prefibrillar oligomers, all of which are desirable characteristics of a robust amyloid detection tool. However, this interaction has only been reported using data from fluorescence microscopy, which is more labor- and time-intensive than the simple bulk fluorescence measurements that can be applied when using amyloid dyes whose fluorescence properties are affected upon binding. To this end, we have developed an FP-based system for amyloid detection utilizing the principle of Förster resonance energy transfer (FRET) that exploits the inherent amyloid affinity of FPs while allowing for simple detection through spectroscopic methods.

The PAPf39 amyloid system was used as the model for this work, as superfolder GFP (sfGFP) has been previously determined to bind these fibrils with high affinity (58). Amyloid aggregates formed by the PAPf39 peptide are found naturally in human semen and have been found to dramatically enhance human immunodeficiency virus (HIV) infection by mediating the viral particle-host cell interaction (33; 52).

The probe we have designed is comprised of two new FRET-compatible sfGFP derivatives separated by a globular linker domain with flexible hinge regions. Here, we demonstrate the ability of this sensor molecule to report the presence of PAPf39 fibrils through changes in FRET, while remaining insensitive to the presence of monomeric peptide and amorphous protein aggregates. This work serves as proof of concept for further development of an FP-based sensor platform for amyloid detection and quantification.

## 2. Results and Discussion

We have previously established, using confocal microscopy, that sfGFP binds the core of various amyloid fibrils (58). However, the binding does not change the fluorescence profile of the protein, suggesting that there are very small, if any, perturbations in the 3D structure. It also indicates that surface residues of GFP are directly involved in binding. We, therefore, set out to develop a system consisting of two FPs that exhibit FRET-compatible fluorescence properties (Figure 1). In this system, two FRET-compatible FP molecules are tethered by a flexible linker such that binding to the fibril surface induces a change in FRET efficiency between the donor and acceptor FP domains that can be observed through changes in fluorescence upon donor excitation. Such a system needs to satisfy the following conditions: 1. The surfaces of the two FPs must be identical so that their intrinsic binding mechanism to the amyloid fibril is the same; 2. The emission spectrum of one FP must have significant overlap with the excitation spectrum of the other to allow for FRET; 3. Both FPs should exhibit relatively high brightness and photostability, as these are generally desirable FP characteristics. To this end, we decided to alter sfGFP, which has high thermostability, to derive proteins with the fluorescence properties of a well-established FRET pair consisting of the cyan fluorescent protein, Aquamarine (11), and yellow fluorescent protein, mCitrine (16), while retaining the surface identical to sfGFP. GFP derivatives of different colors have been developed mainly through a high-throughput screening following error-prone PCR of the gene encoding this protein. As a result, while many different FP with different fluorescence properties were discovered in this way, they contain mutations that are not directly related to the structure and properties of the chromophore (34). In particular, the mutations on the surface of the protein and away from the chromophore are very unlikely to be relevant to the final fluorescence profile. We thus compared the sequence and structure of sfGFP with those of Aquamarine and mCitrine to identify the residues directly affecting the structure and environment of the chromophore. There is a difference in 9 residues between sfGFP and Aquamarine (Figure 2A). These residues were examined on the crystal structure of sfGFP (PDB ID 2B3P, (37)) to identify the location of the residue side chain; if the residue side chain extends to the interior of the β-barrel, the residue found in Aquamarine was introduced into the sequence of sfGFP. This allowed us to identify five mutations in the vicinity of the chromophore, which were thus retained in the engineered protein termed sfCyan. Similarly, there is a difference in 15 positions between sfGFP and mCitrine (Figure 2B). Of these, nine were retained in the engineered protein termed sfYellow. In both cases, the surface residues of sfGFP were retained. Additionally, the C-terminal nine residues of the sfGFP sequences were not included in sfCyan or sfYellow to simplify the sequences of the fusion constructs. Crystal structures of various GFP constructs have demonstrated the unstructured nature of residues 230-238 [46-50]. Furthermore, truncation studies have shown that removing these residues does not affect the stabilities or fluorescence properties of these proteins [51, 52]. The final sequences of sfCyan and sfYellow proteins as cloned are shown in Figures 2A,B. These proteins were expressed, purified, and characterized by relevant assays. First, we examined the excitation and emission spectra of the resulting individual FP constructs. In Figures 2C,D, the spectral properties of Aquamarine and mCitrine are compared with those of sfCyan and sfYellow, respectively. It is evident that the spectral properties of mCitrine and sfYellow are practically identical, while there is only a minimal difference between the optical properties of Aquamarine and sfCyan. Second, we used confocal microscopy to visualize a mixture of our model PAPf39 amyloid fibrils with equimolar concentrations of sfCyan and sfYellow. This experiment demonstrated the independent abilities of sfCyan and sfYellow to bind PAPf39 fibrils (Figure S1), thus supporting the initial goal of changing the color of the FP while maintaining the ability to bind fibrils.

**Figure 1:**
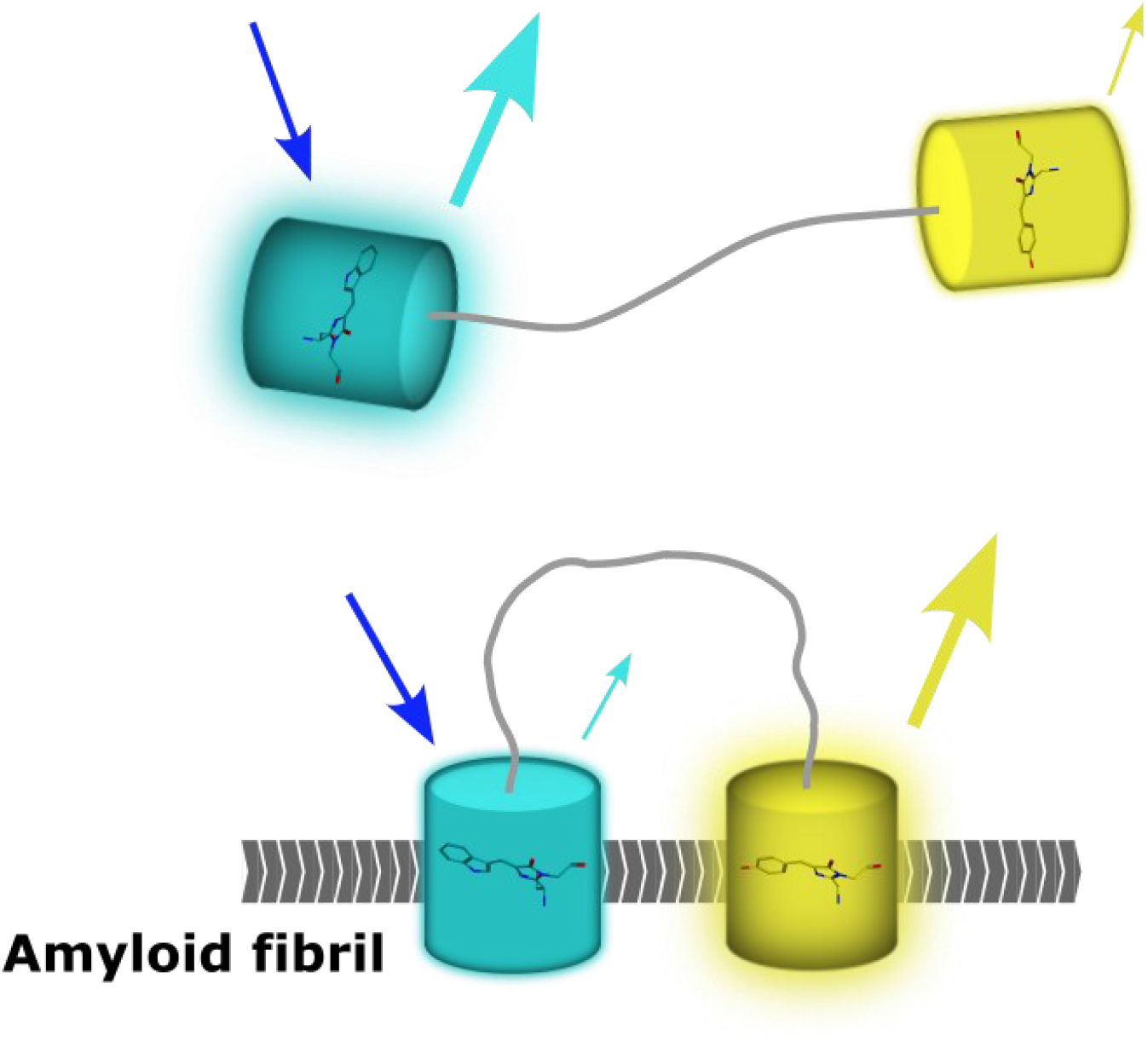
Concept of FRET-based reporter for fluorescent protein-PAPf39 fibril binding. Donor fluorescent protein is shown in cyan, and acceptor fluorescent protein is shown in yellow. Upon excitation of the donor, where the FP domains are far apart in solution, there is little energy transfer to the acceptor. In the fibril-bound state, the FP domains come closer together, resulting in an increase in energy transfer that can be observed through changes in the total emission spectrum.

**Figure 2:**
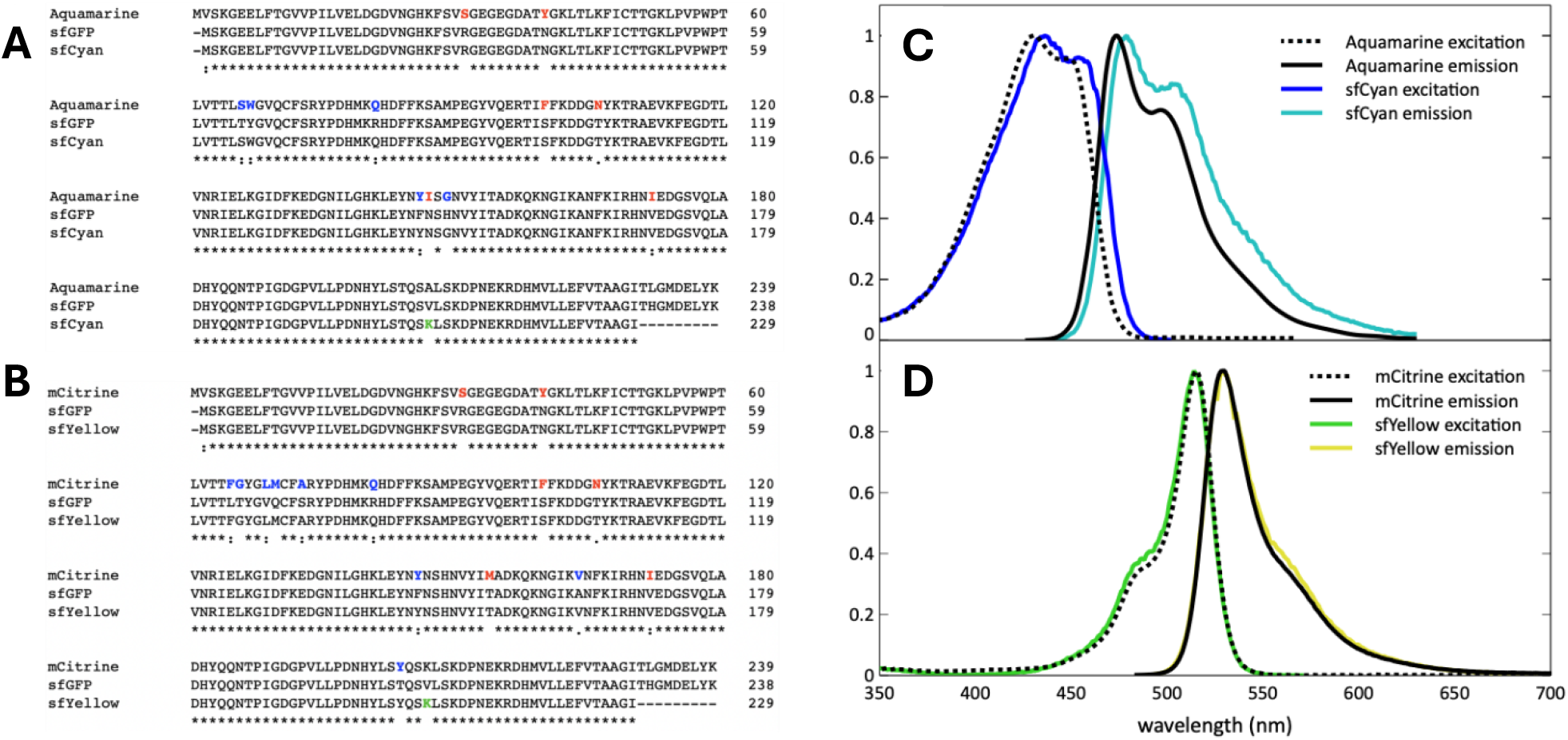
Fluorescent protein sequence alignments and excitation and emission spectra of sfCyan and sfYellow. (A) Alignment of mCitrine (1-230), sfGFP (1-229), and sfYellow sequences. (B) Alignment of Aquamarine (1-230), sfGFP (1-229), and sfCyan sequences. Multiple sequence alignments were generated using Clustal Omega (v1.2.4, (48)). Residues colored in red denote residues that were mutated to the residue found at that position of sfGFP. Residues colored in blue denote residues that were kept from the base fluorescent protein. (C) Aquamarine excitation (dotted black) and emission (dashed black) spectra from literature (11). sfCyan excitation (blue) and emission (cyan) spectra. (D) mCitrine excitation (dotted black) and emission (dashed black) spectra from literature (16). sfYellow excitation (green) and emission (yellow) spectra. Spectra are normalized to the peak intensity.

After initial characterization of the engineered sfCyan and sfYellow proteins, we proceeded to generate a covalent heterodimer of these two proteins connected by a linker to function as a FRET sensor. We used a SpyTag-SpyCatcher system (59) that provides several essential features for making heterodimers of sfCyan and sfYellow. First, SpyTag and SpyCatcher form a covalent isopeptide bond, ensuring stable and durable linkage between two proteins expressed independently. Second, the reaction is rapid and high yield over a broad range of conditions, making it highly efficient and adaptable for forming covalent heterodimers. Third, individual proteins fused to SpyTag or SpyCatcher are genetically encoded, thus providing modularity to the system for possible downstream optimization efforts.

We used the minimal sequences of the SpyTag and SpyCatcher components required for efficient heterodimerization, SpyTag003 (ST) and ΔN1ΔC2SpyCatcher (SC) (27; 22). A decapeptide with the sequence <u>DVGSSGSSLQ</u> was used as a linker (Figure 3A) between SpyTag and sfCyan (SpyTag at C -terminus or N-terminus of sfCyan termed STCyan and CyanST, respectively) and between sfYellow and SpyCatcher (termed YellowSC). Combining two different SpyTag-fusions of sfCyan with YellowSC generates two alternative architectures for relative positions of FP pairs – “parallel” (YellowSC-CyanST) and “antiparallel” (YellowSC-STCyan) – where the FP domains exist either on the same face or opposite faces of the SpyTag/SpyCatcher globular heterodimer (Figure 3A). SDS-PAGE demonstrates near complete covalent heterodimerization of the STCyan and YellowSC constructs following 20 minutes of co-incubation (Figure 3B). Confocal microscopy of YellowSC-STCyan and YellowSC-CyanST constructs incubated with PAPf39 fibrils confirmed the ability of these molecules to bind the PAPf39 amyloid fibril (Figure 3C). To confirm the presence of FRET in the fibril-bound state, FRET acceptor-photobleaching microscopy was performed using samples of YellowSC-STCyan incubated with PAPf39 fibrils. Briefly, this method involves bleaching the acceptor fluorophores in a region of interest using intense excitation and measuring the fluorescence intensity of the donor before and after bleaching. This photobleaching of acceptor fluorophores results in an increase in donor fluorescence intensity directly related to the efficiency of energy transfer. This, in turn, allows for the direct calculation of FRET efficiency within a region of interest. These experiments confirmed the presence of energy transfer between sfCyan and sfYellow within the YellowSC-STCyan construct when bound to PAPf39 fibrils with a FRET efficiency of 0.40 ± 0.05 (Figure S2). Then, using Equations 5 and 6, we can estimate the average distance between sfCyan and sfYellow fluorophores, and, hence, the average distance between adjacent FP binding sites on the PAPf39 fibril, of 53 ± 3 Å. Identical experiments conducted using the YellowSC-CyanST construct yielded a bound-state FRET efficiency of 0.48 ± 0.04, corresponding to FP binding site spacings of 49 ± 2 Å. The similarity of bound-state FRET efficiency measurements and associated FP binding site spacings obtained for both constructs (despite exhibiting significantly different free-state FRET efficiencies, see Figure 4B) further support the estimates of spacing between FP-binding sites on the PAPf39 fibril.

**Figure 3:**
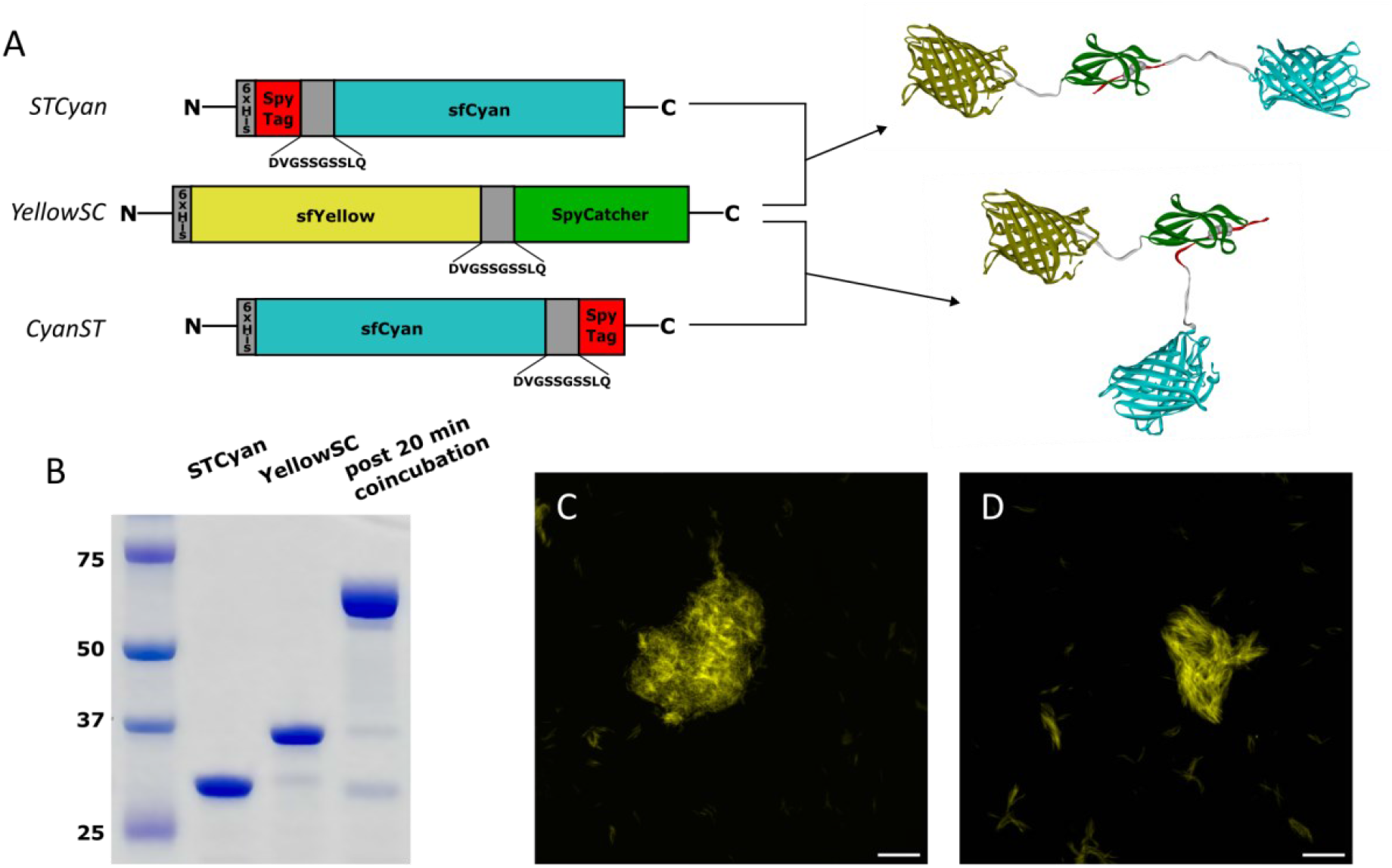
YellowSC-STCyan and YellowSC-CyanST construct structures and validation of PAPf39 amyloid-binding. (A) Fusion constructs STCyan, CyanST, and YellowSC and ribbon models of YellowSC-STCyan (“antiparallel”) and YellowSC-CyanST (“parallel”). Isopeptide bond in SpyTag003-ΔN1ΔC2SpyCatcher dimer shown in CPK representation. (B) Coomassie-stained SDS-PAGE gel demonstrating covalent heterodimerization of STCyan and YellowSC after 20 min incubation. Molecular mass markers in kDa. (C and D) Confocal microscopy image of 30 μM PAPf39 fibrils incubated with 1 μM YellowSC-STCyan (C) or 1 μM YellowSC-CyanST (D). Image acquired with the 405 nm argon laser and 525-570 nm bandpass emission filter. The scale bar is 20 μm.

**Figure 4:**
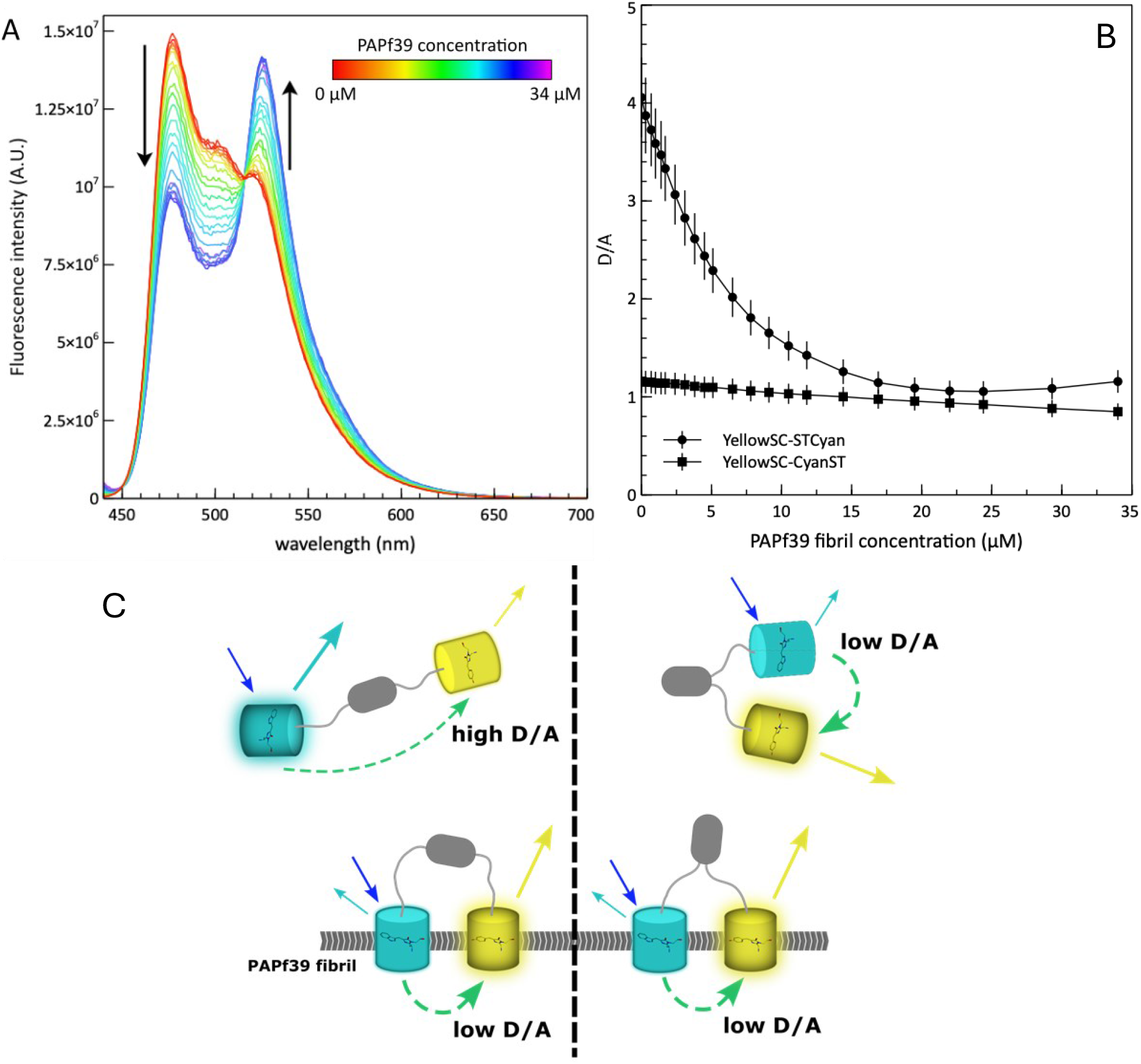
YellowSC-STCyan emission spectrum shows PAPf39 fibril concentration dependence. (A) Raw YellowSC-STCyan emission spectra from 440-700 nm (*λ*_ex_ = 430 nm) as a function of PAPf39 fibril concentration. Arrows indicate simultaneous intensity decrease at 477 nm and increase at 525 nm, corresponding to the emission maxima of sfCyan and sfYellow, respectively. (B) Average D/A of YellowSC-STCyan (circles) and YellowSC-CyanST (squares) derived from raw emission spectra over three replicates (see supplementary Figure S7). Error bars represent 10% error. (C) Model to explain the observed difference in D/A sensitivity to PAPf39 fibrils between YellowSC-STCyan (left) and YellowSC-CyanST (right).

To quantitatively assess the amyloid sensing capabilities of the YellowSC-STCyan and YellowSC-CyanST constructs in bulk solution using conventional fluorescence spectroscopy, we measured the emission spectrum in the presence of increasing concentrations of preformed PAPf39 fibril. The raw emission spectra of the YellowSC-STCyan construct as a function of PAPf39 fibril concentration are shown in Figure 4A. It is evident that as the concentration of fibril increases, there is a simultaneous intensity decrease at 477 nm (emission maximum of sfCyan) and an increase of intensity at 525 nm (emission maximum of sfYellow); these changes correspond to an increase in the degree of energy transfer, indicative of a decrease in the average distance between FP domains in the sensor construct, upon binding the fibril surface. To quantify the emission intensity changes of each FP, the observed emission spectrum at each fibril concentration was analyzed to extract a derived parameter, D/A, as a proxy for FRET efficiency. D/A is the ratio of the sfCyan and sfYellow reference spectrum scaling parameters upon linear unmixing (see Materials and Methods and Equation 4).

Figure 4B shows the changes in D/A for both YellowSC-STCyan and YellowSC-CyanST as functions of PAPf39 fibril concentration in monomer equivalents. The degree of change in D/A of the YellowSC-STCyan construct is more pronounced: it starts at high D/A and follows a hyperbolic trend expected for equilibrium binding processes (see also Figure S3 for reproducibility). However, the degree of change in D/A of the YellowSC-CyanST construct is minimal: it starts at a low D/A and changes less than 1 D/A unit over the course of the titration. This difference observed between these two constructs can be attributed to the structural differences that ultimately result in different degrees of energy transfer in the free state. The initial D/A value (i.e. D/A in the absence of fibrils) provides information on the degree of energy transfer in the free state. It demonstrates that the “parallel” YellowSC-CyanST construct exhibits a much higher degree of energy transfer than the “antiparallel” YellowSC-STCyan. Taken together with the fact that the bound-state FRET efficiency values and the D/A at the end of the titrations are relatively similar, it can be deduced that the lower amyloid sensitivity observed using YellowSC-CyanST relative to the antiparallel construct is likely due to the fact that, although the average distance between FP domains in the bound-state is similar across both constructs, the average distance between FP domains does not undergo as significant a change in average distance from the free-to bound-state (Figure 4C).

To demonstrate that the observed changes in D/A are due to a specific interaction with the PAPf39 fibril, similar experiments were conducted using the YellowSC-STCyan construct and monomeric PAPf39; these data showed no change in D/A over the same concentration range, supporting the notion that the change in D/A in the YellowSC-STCyan construct is amyloid-specific and reports on the fibril binding event (Figure S4A). Additionally, amorphous aggregates made from heat-denatured bovine serum albumin (BSA) also do not induce a change in YellowSC-STCyan D/A over an analogous concentration range, showing that the FRET signal is not affected by the presence of protein aggregates that do not interact with FPs (Figure S4B) in agreement with our previous assessment (58).

Finally, to address the question of potential non-specific irreversible adsorption of FPs to fibrils in general and of the YellowSC-STCyan construct in particular, we performed competition titrations of PAPf39 fibrils with a sfGFP variant that does not show fluorescence in the same wavelength region as the YellowSC-STCyan emission spectrum. This variant contains the Y66F mutation, which results in a significant blue-shift in the excitation and emission spectra (*λ*_ex,max_=356 nm, *λ*_em,max_=428 nm), as well as a significant decrease in quantum yield (QY = 0.013) (17; 35; 38; 26). We termed this sfGFP variant “dark” sfGFP (dsfGFP), as it is non-fluorescent with the sets of experimental parameters used to probe YellowSC-STCyan. The dsfGFP variant is a properly folded protein adopting a β-barrel structure (4) that retains the same surface-presenting sequence as both sfCyan and sfYellow, allowing it to compete for the same PAPf39 fibril binding sites without contributing to D/A. Figure 5 shows the decrease in D/A upon titration of YellowSC-STCyan with PAPf39 fibril until D/A values started to reach a plateau. At that point, the addition of dsfGFP results in the recovery of D/A, indicative of successful competition of YellowSC-STCyan from the fibril, suggesting both the reversibility and specificity of the YellowSC-STCyan interaction with PAPf39 fibrils.

**Figure 5:**
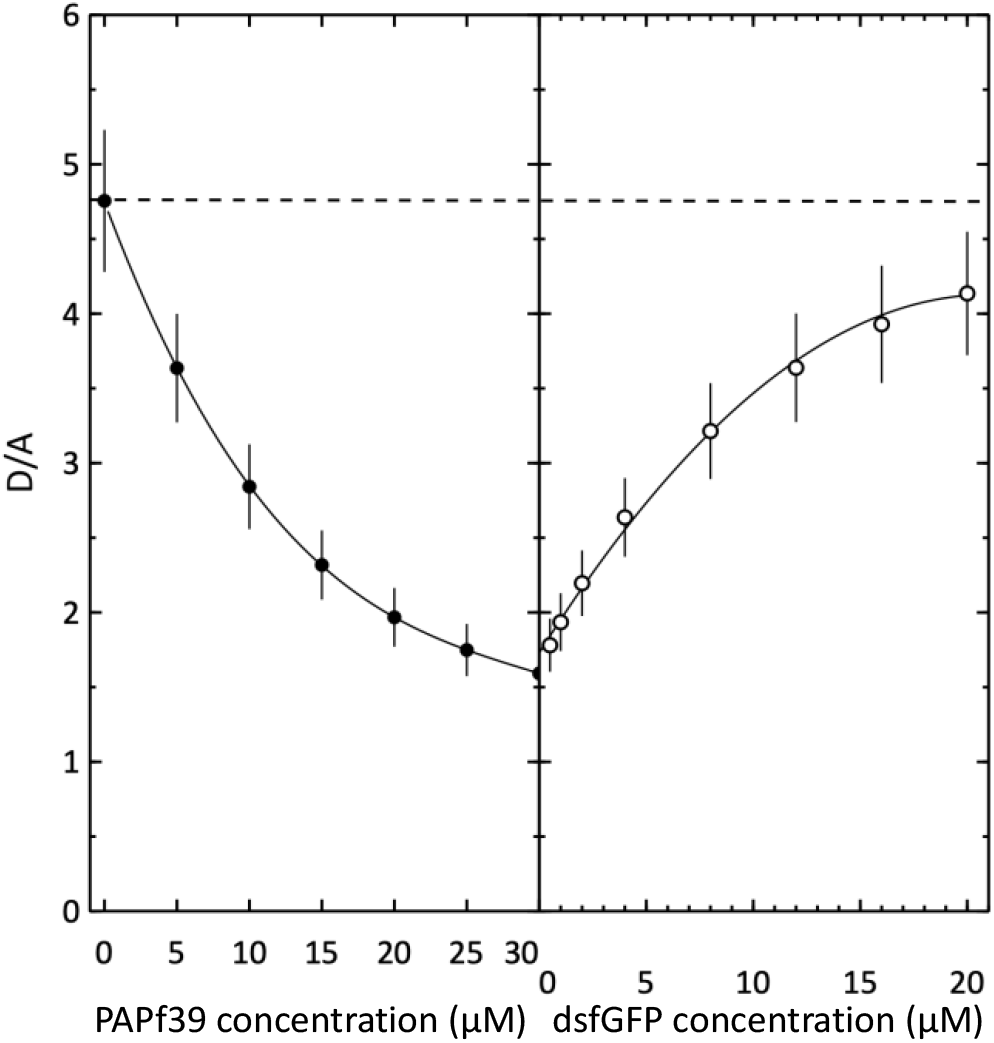
YellowSC-STCyan binding to PAPf39 fibrils is reversible and specific. PAPf39 fibrils titrated into 1 μM YellowSC-STCyan up to 30 μM fibril (left panel, solid symbols). (right) YellowSC-STCyan D/A (right panel, open symbols) recovers towards basal D/A (dotted line) as a function of dsfGFP concentration. Error bars represent 10% error.

## 4. Conclusions

The bulk solution sensitivity of the YellowSC-STCyan molecule to the presence of PAPf39 fibrils and lack of response to equivalent concentrations of either monomeric peptide or amorphous protein aggregates supports the concept of using an FP-based FRET system for detecting amyloid fibrils. These data, along with the confirmation that the observed change in D/A is a result of a specific, reversible interaction with the fibrils, corroborate the previously published work illustrating the FP-fibril interaction (58). The foundational work in reporting the FP-amyloid interaction was conducted primarily using fluorescence microscopy, as most other classic biophysical methods of identifying protein-protein interactions are not amenable to the study of insoluble material, including amyloid aggregates, and because the FP fluorescence properties are insensitive to the interaction. Through exploiting the principle of FRET, we have engineered a fluorescence-based sensor that has made the FP-fibril interaction observable through bulk fluorescence measurements that can be used to yield more quantitative information regarding the FP-fibril interaction than what is feasible to achieve using fluorescence microscopy data. This work was completed using a model amyloid system, PAPf39, but, given the generalizability of the FP-amyloid interaction across numerous systems (58), this can be extended to assess the ability of our sensor platform to report on the presence of these other fibrils, as well.

## 4. Materials and Methods

### Cloning of protein constructs

The fluorescent protein and heterodimerization domains are linked through decapeptide with AatII and PstI restriction site scars on respective termini (-DVGSSGSSLQ-). Plasmids including 6xHis-sfCyan, 6xHis-sfYellow, 6xHis-SpyTag003-DVGSSGSSLQ-sfCyan (STCyan), 6xHis-sfCyan-DVGSSGSSLQ-SpyTag003 (CyanST), 6xHis-sfYellow-DVGSSGSSLQ-ΔN1ΔC2SpyCatcher (YellowSC), and 6xHis-dsfGFP cloned into the pGia expression vector between the SphI and HindIII restriction sites were obtained from Twist Bioscience (South San Francisco, CA). Sequences are listed in the Supplementary Table S1.

### Expression and purification of fluorescent proteins

*E. coli* BL21(DE3) cells containing the expression plasmid were grown overnight in a 5 mL starter culture of 2xYT media containing 100 μg/mL ampicillin at 37°C. 1 mL starter culture was pelleted at 1,800 x g and resuspended in fresh media, which was used to inoculate 1 L of fresh media in a baffled Fernbach flask. The culture was then incubated at 37°C while shaking at 225 rpm until OD_600_ 0.6-0.8 where expression was induced using 1 mM IPTG. The induced culture was incubated at 37°C while shaking at 225 rpm for 4-8 hr. Cells were harvested by centrifugation at 2,900 x g and 4°C for 30 min, spent media was discarded, and pellets were frozen at -20°C for >12 hr. The cell pellets were then thawed at room temperature, resuspended with ∼10 mL 20 mM Tris pH 8.0, 200 mM NaCl, 10 mM imidazole and lysed by passing twice through an ice-cold French pressure cell. Cell debris was pelleted by centrifugation of the cell lysate at 13,600 x g and 4°C for 1 hr. The supernatant was loaded to a ∼30 mL Ni^2+^-NTA column (ThermoFisher Scientific), washed with 20 mM Tris pH 8.0, 200 mM NaCl, 10 mM imidazole, and eluted using 20 mM Tris pH 8.0, 200 mM NaCl, 200 mM imidazole. For covalent dimerization of ST/SC fusions, the concentrations of both ST/SC pairs were estimated by measuring the absorbance of each eluant at their respective excitation maxima (sfCyan, 430 nm; sfYellow, 515 nm). The constructs were mixed at an approximately equimolar ratio and incubated at 25°C with shaking at 225 rpm for 20 min to allow for the ST/SC dimerization reaction to occur.

The nickel column eluant was then dialyzed against 100 mM Tris pH 8.0, 5 mM EDTA using a 3.5 kDa MWCO dialysis membrane (Fisher Scientific) at 4°C overnight. The sample was loaded to a ∼30 mL DEAE Sepharose G25 Fast Flow column (Cytiva Life Sciences, Marlborough, MA) and eluted using a linear 0-500 mM NaCl gradient in 100 mM Tris pH 8.0, 5 mM EDTA, followed by size exclusion chromatography using a 2.5 x 120 cm Sephadex G-75 column (Cytiva Life Sciences) in PBS pH 7.4 with 0.02% sodium azide (will be referred to as PBSa). Fractions were checked for purity via SDS-PAGE using SurePAGE Bis-Tris, 8-16% acrylamide precast gels and Tris-MES buffer (GenScript, Piscataway, NJ), as well as MALDI-TOF. Pure fractions were pooled, concentrated using a 10 kDa MWCO centrifugal filter (Pall Corporation, Port Washington, NY), and stored at -20°C in PBSa and 20% glycerol.

### PAPf39 expression and purification

The recombinant expression and purification of the PAPf39 peptide was done as described previously (46). The pTXB1 expression vector containing the fusion expression construct, 6XHis-Ubiquitin-PAPf39-Intein, was transformed into *E. coli* BL21(DE3) cells using standard heat shock transformation protocol. Cells were grown overnight in a 10 mL starter culture of 2xYT media with 100 μg/mL ampicillin. 1.5 mL starter culture was used to inoculate 1 L 2xYT media containing 100 μg/mL ampicillin, which was incubated at 30°C while shaking at 225 rpm until OD_600_ reached ∼0.4. Expression was induced using 1 mM IPTG and the induced cultures were incubated at 22°C while shaking at 225 rpm for 4 hr. Cells were harvested and lysed, and the resulting lysate containing 6XHis-Ubiquitin-PAPf39-Intein was purified through Ni^2+^-NTA affinity chromatography following the same protocols as described for the fluorescent protein constructs.

For simultaneous intein self-cleavage and TEV protease cleavage of the ubiquitin tag, 1 mg TEV protease was added for every 50 mg 6XHis-Ubiquitin-PAPf39-Intein. Additionally, for every 100 mL of solution, 1 mL 500 mM EDTA pH 8.0, 800 μL βME, and 200 μL 1 M PMSF were added. The TEV protease reaction was incubated for ∼24 hr at 4°C with constant stirring. The TEV protease cleavage reaction was loaded to a ∼30 mL CM Sepharose Fast Flow column (Cytiva Life Sciences) in 20 mM Tris, 5 mM EDTA pH 8.0 and eluted using a 0-1 M ammonium acetate gradient in 20 mM Tris, 5 mM EDTA pH 8.0. Fractions containing PAPf39 were pooled and loaded to a Discovery Bio Wide Pore C18 column (25 cm x 10 mm, Supelco Sigma-Aldrich, Bellefonte, PA) in 0.05% trifluoroacetic acid and eluted using a 0-100% methanol gradient. Pure fractions were pooled and lyophilized, and the dry peptide was stored at -20°C.

### Measuring protein and peptide concentrations

All protein and peptide concentrations were measured by absorbance at 280 nm in a quartz cuvette using a UV-Vis spectrophotometer (Hitachi U-2900, Tokyo, Japan). Extinction coefficients and molecular weights were calculated using the ExPASy Protein Parameters server (14).

### Matrix-assisted laser desorption/ionization-time-of-flight (MALDI-TOF)

MALDI-TOF mass spectrometry was used to confirm the identities and purity of all proteins and peptides used. MALDI-TOF was performed using a Bruker Ultraflex III MALDI TOF/TOF instrument with the flexControl and flexAnalysis software (Bruker Daltonics, Billerica, MA) for data acquisition and analysis, respectively. PAPf39 peptide was analyzed using an α-cyano-4-hydroxycinnamic acid (CHCA) matrix. All other proteins were analyzed using a matrix comprised of a mixture of CHCA and 2,5-dihydroxybenzoic acid (DHB) (49).

### Sedimentation equilibrium analytical ultracentrifugation

Samples of YellowSC-STCyan in PBSa were loaded into 2-sector cells and centrifuged using a ProteomeLab XL-A analytical ultracentrifuge equipped with an An-60 Ti 4-cell rotor (Beckman Coulter, Brea, CA) at 23°C. Absorbance profiles at 430 nm were recorded every 6 hr. Samples were centrifuged consecutively at 15,000 rpm, 18,000 rpm, and 22,000 rpm after achieving equilibration for ≥18 hr at each speed.

To estimate the apparent molecular weight, equilibrium absorbance scans at each speed were globally fit to the following single-component exponential model using nonlinear regression NLREG as described (50):

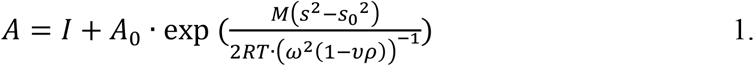

where *M* is the apparent molecular weight, *A* is the absorbance at 430 nm as a function of radius, *I* is the baseline absorbance offset at 430 nm, *A*_*0*_ is the absorbance at 430 nm at an arbitrary reference radius, *s* is the radius along the column in cm, *s*_*0*_ is an arbitrary reference radius, *ω* is the angular velocity of the rotor in rad/s, *υ* is the partial molar volume of the protein in mL/mol, *R* is the gas constant (8.3145 x 10^7^ g cm^2^ mol^-1^ K^-1^), *T* is the temperature in K, and *ρ* is the solvent density in g/mL (assumed to be equal to that of water, ∼1 g/mL). The partial molar volume of YellowSC-STCyan was calculated to be 0.738 mL/mol from the summation of experimentally determined partial molar volumes at 25°C of the constituent amino acids, peptide bonds, and N- and C-termini, and the calculated molecular weight of the protein (30). The analysis supported the monomeric nature of the purified YellowSC-STCyan construct, with the estimated molecular mass to be 63.8 ± 1.9 kDa, close to the molecular mass of 65.9 kDa calculated from the amino acid sequence (Figure S5).

### Quantum yield measurement

The fluorescence quantum yields of sfYellow and sfCyan at 10°C, 25°C, and 40°C were determined using the comparative method, where the proportionality of fluorescence versus absorbance for the sample of unknown quantum yield is compared to that of a reference sample with well-documented quantum yield (36; 55). This quantum yield determination method is described by the following equation:

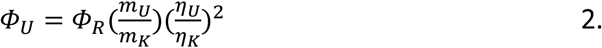

where U represents the unknown sample, K represents the reference sample of known quantum yield, Φ represents the quantum yield, m represents the slope of the plot of integrated fluorescence intensity versus absorbance, and *η* represents the solvent refractive index.

Rhodamine 6G (ThermoFisher Scientific) in 100% ethanol at 25 °C was used as the reference sample for sfYellow quantum yield measurement (Φ = 0.95), and ATTO-425 (Sigma-Aldrich, Bellefonte, PA) in PBS pH 7.4 at 25°C was used as the reference for sfCyan (Φ = 0.9) (25). sfCyan and sfYellow quantum yields were measured in PBSa. Solvent refractive indices at the respective temperatures were measured in triplicate using the Bausch & Lomb (Laval, Canada) Abbe-3L (33-45-58) refractometer, with the temperature regulated using a NESLAB Instruments (Newington, NH) RTE100-NonCFC water bath. 5 independent solutions of each sample were prepared with different absorbance values less than 0.1 at the excitation wavelength (sfYellow/rhodamine 6G: 485 nm; sfCyan/ATTO-425: 430 nm). Absorbance spectra were recorded using a quartz microcuvette with a 10 mm pathlength in a U-2900 UV-Vis spectrophotometer thermoregulated with a Wavelength Electronics (Bozeman, MT) LFI-3751 Temperature Controller; emission spectra were recorded in the same cuvette in a FluoroMax-4 spectrofluorometer (Horiba Jobin Yvon, Kyoto, Japan) thermoregulated with a NESLAB Instruments RTE100-NonCFC water bath. The absorbance at the excitation wavelength was corrected for background absorbance and baseline drift, and the emission spectra were corrected for excitation light intensity and background signal. The integrated fluorescence intensity was calculated via trapezoidal approximation from 490-700 nm for sfYellow/rhodamine 6G and from 435-675 nm for sfCyan/ATTO-425, then plotted against the absorbance at the excitation wavelength with the y-intercept set to zero.

### PAPf39 fibril formation

Dry peptide was dissolved in ice-cold 3.5 mM HCl and centrifuged at 21,900 x g and 4°C for 20 min to pellet any insoluble particles. The clarified peptide solution was transferred into a fresh tube on ice, and the concentration was measured by absorbance at 280 nm using an extinction coefficient of 0.65 mL mg^-1^ cm^-1^. The concentration was adjusted to 6 mg/mL using 3.5 mM HCl and diluted to 2 mg/mL using cold 1.5X-concentrated PBS pH 8.1. The solution was immediately immersed in a gyratory water bath shaker (37°C and 180 rpm) for ∼48 hr. Fibril concentrations are reported in units of monomers.

### BSA amorphous aggregate formation

2 mg/mL BSA solution was prepared by weighing out dry BSA (VWR) and dissolving in the appropriate volume of PBS. The sample was boiled for 20 min and left at room temperature to cool.

### Atomic force microscopy (AFM)

AFM was used to confirm the presence of fibrillar structures in all preformed fibril samples (data not shown). Samples were deposited on freshly cleaved mica (SPI Supplies, Westchester, PA) glued to a microscope slide and air-dried before rinsing 3 times with 1 mL milli-Q water. The sample was left to dry entirely before imaging on an MFP-3D atomic force microscope (Asylum Research, Santa Barbara, CA). Probes (Catalog #AC240TS-R3, Oxford Instruments, Abingdon, England) used for imaging have an aluminum-coated cantilever and a non-coated tip with a radius of 7 nm, resonant frequency of 70 kHz, and spring constant of 2 N/m. Images were collected and processed using the Igor Pro MFP3D (Wavemetrics Inc., Portland, OR) software.

### Confocal fluorescence microscopy

Fluorescence imaging was conducted using the Leica (Wetzlar, Germany) TCS SP8 3X STED microscope and the LAS X software. Slides (75 mm x 25 mm x 1 mm, VWR) and coverslips (22 mm x 22 mm, No. 1.5, VWR) were cleaned by soaking in 30% ethanol and dried using a compressed air stream. 10 μL of the sample was spotted on the coverslip, placed on the slide, and the edges of the coverslip were sealed with clear nail polish. For imaging YellowSC-CyanST and YellowSC-STCyan sfYellow in the presence of PAPf39 fibrils, 1 μM heterodimeric FP molecule was co-incubated with 30 μM PAPf39 fibril and allowed to equilibrate for ∼20 min at room temperature before imaging. The microscope was equipped with a 63X objective (Leica HC PL APO, Oil immersion, 1.40 N.A.) and the 514 nm argon laser, and emission was detected from 525-570 nm. For imaging untethered sfCyan and sfYellow in the presence of PAPf39 fibrils, 5 μM each of sfCyan and sfYellow were incubated with 90 μg/mL PAPf39 fibril and allowed to equilibrate for ∼20 min at room temperature before imaging. The microscope was equipped with a 100X objective (Leica HC PL APO, Oil immersion, 1.4 N.A.); sfCyan was excited using the 405 nm argon laser, and emission was detected from 420-490 nm, while sfYellow was excited using the 514 nm argon laser, and emission was detected from 525-570 nm. The pinhole for all sets of acquisition parameters was set to 1 Airy unit.

### FRET acceptor photobleaching microscopy

Triplicate experiments were performed in using a Leica TCS SP8 3X STED microscope equipped with a 63X objective (Leica HC PL APO, Oil immersion, 1.40 N.A.) and using the FRET-AB module of the LAS X software. Samples were prepared by mixing the components in a 1.5 mL microcentrifuge tube to final concentrations of 1 μM YellowSC-STCyan or YellowSC-CyanST and 30 μM PAPf39 fibril and allowed to equilibrate for ∼20 min at room temperature. Slides were cleaned by soaking in 30% ethanol and dried using a compressed air stream. 10 μL of the sample was spotted on the coverslip, placed on the slide, and the edges of the coverslip were sealed with clear nail polish. 184.9 μm x 184.9 μm images corresponding to direct excitation and emission of both sfCyan and sfYellow were acquired pre- and post-bleaching of sfYellow within a defined region of interest. sfCyan and sfYellow were imaged with argon laser lines at 458 nm (emission detected from 470-500 nm) and 514 nm (emission detected from 525-570 nm), respectively. The pinhole was set to 1 Airy unit for all sets of acquisition parameters. sfYellow was bleached to <10% of the pre-bleaching intensity using the 514 nm Argon laser for 15 frames. The AccPbFRET plugin (41) for Fiji (44) was used to calculate the pixel-by-pixel FRET efficiency utilizing the following equation:

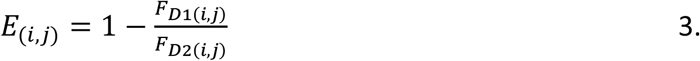

where E represents FRET efficiency at a given pixel (i,j), and F_D1_ and F_D2_ represent the donor fluorescence intensities before (1) and after (2) acceptor photobleaching. Background intensity was subtracted during this analysis, and pixels were registered to account for shifts within the XY-plane. A histogram of FRET efficiencies at each pixel (0.36 μm x 0.36 μm) within the sfYellow-depleted region of interest was generated and fit using NLREG (47) to a simplified normal distribution with the following form:

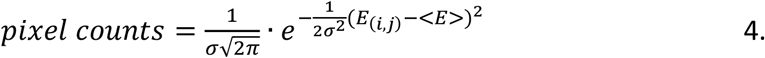

where *<E>* represents the mean FRET efficiency within the region of interest and σ represents the standard deviation of the distribution. To estimate the distance between sfCyan and sfYellow fluorophores in the fibril-bound state from *<E>* measurements, the method described by Vogel, et al. (54) was employed. The widely used assumption of a dynamic isotropic regime (i.e. κ^2^ = 2/3) in the calculation of the Förster distance (R_0_) is violated in cases where the fluorophores rotate much slower than their excited state lifetimes, as is the case with FP FRET pairs. The relationship between the distance between fluorophores (R) and *<E>* of FRET pairs within the static isotropic regime (ϕ) was empirically determined and tabulated to allow for more accurate distance estimations. The corresponding ϕ of each *<E>* value was recorded and used to calculate the distance between fluorophores using the following equations:

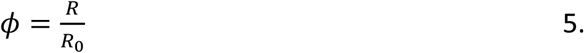

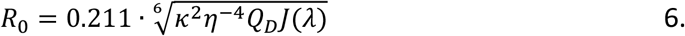

where *R*_*0*_ represents the Förster radius of the sfCyan/sfYellow FRET pair, calculated to be 53.78 Å from the solution refractive index (*η*), quantum yield of sfCyan (Q_D_), and spectral overlap integral, J(*λ*), measured at 23°C (57; 16; 11) (orientation factor, κ^2^, assumed to be 0.667). The distance estimates (YellowSC-STCyan, n=34; YellowSC-CyanST, n=30) were then averaged to determine the distance between FP-binding sites on the PAPf39 fibril.

### Fluorescence spectroscopy fibril titrations

Triplicate experiments were performed on a Fluoromax-4 spectrofluorometer (Horiba Jobin Yvon, Kyoto, Japan) at 23°C using a quartz microcuvette with a 10 mm pathlength. For the titration experiments, 1 μM of YellowSC-STCyan or YellowSC-STCyan in PBS was titrated with a stock sample of preformed PAPf39 fibrils, monomeric PAPf39, or preformed amorphous BSA aggregate supplemented with the FP sensor to maintain a constant FP concentration. After each titration point, the sample was allowed to equilibrate for 1 min under continuous stirring. The sample was then excited at 430 nm (1.5 nm slit widths), and the emission spectrum scaled by wavelength-dependent lamp intensity (i.e., S/R) was detected from 440-700 nm (2 nm slit widths) with 90 angle geometry. Inner filter effects resulting from (1) attenuation of excitation light passing through the sample and (2) re-absorbance of emitted photons were assumed not to affect the observed ratios of donor/acceptor fluorescence (i.e., D/A, see below).

The spectra were corrected for direct acceptor excitation by subtracting the emission upon excitation at 430 nm of sfYellow alone from each spectrum. The spectra were then unmixed to separate sfCyan and sfYellow emissions using the following equation:

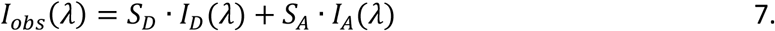

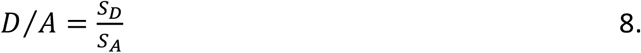

where, at a given wavelength (*λ*), *I*_*obs*_ is the observed fluorescence intensity, *I*_*D*_ and *I*_*A*_ are the reference donor (sfCyan) and acceptor (sfYellow) emission intensities, respectively, that were measured in donor- and acceptor-only samples using the same acquisition parameters, and *S*_*D*_ and *S*_*A*_ are fitted scaling factors applied to the reference donor and acceptor fluorescence intensities such that their sum yields the observed intensity. The qualities of the fits were assessed by monitoring the residuals upon subtracting the reconstructed spectra from the observed spectra. The ratio of *S*_*D*_ to *S*_*A*_ is referred to as D/A.

### Fluorescence spectroscopy dsfGFP competition experiments

Triplicate experiments were performed on a Fluoromax-4 spectrofluorometer at 23°C using a quartz microcuvette with a 10 mm path length. 1 μM of YellowSC-STCyan in PBSa was titrated with a stock sample of preformed PAPf39 fibrils containing 1 μM of YellowSC-STCyan to maintain a constant fluorescent protein concentration. Once the PAPf39 fibril concentration reached 30 μM, the sample was titrated with a stock sample of dsfGFP containing 1 μM YellowSC-STCyan and 30 μM PAPf39 fibril. After each titration point, the sample was allowed to equilibrate for 1 min under constant stirring, and the YellowSC-STCyan emission spectrum was recorded and unmixed using the same procedure described above.

## Supplementary Information

**Figure S1.**
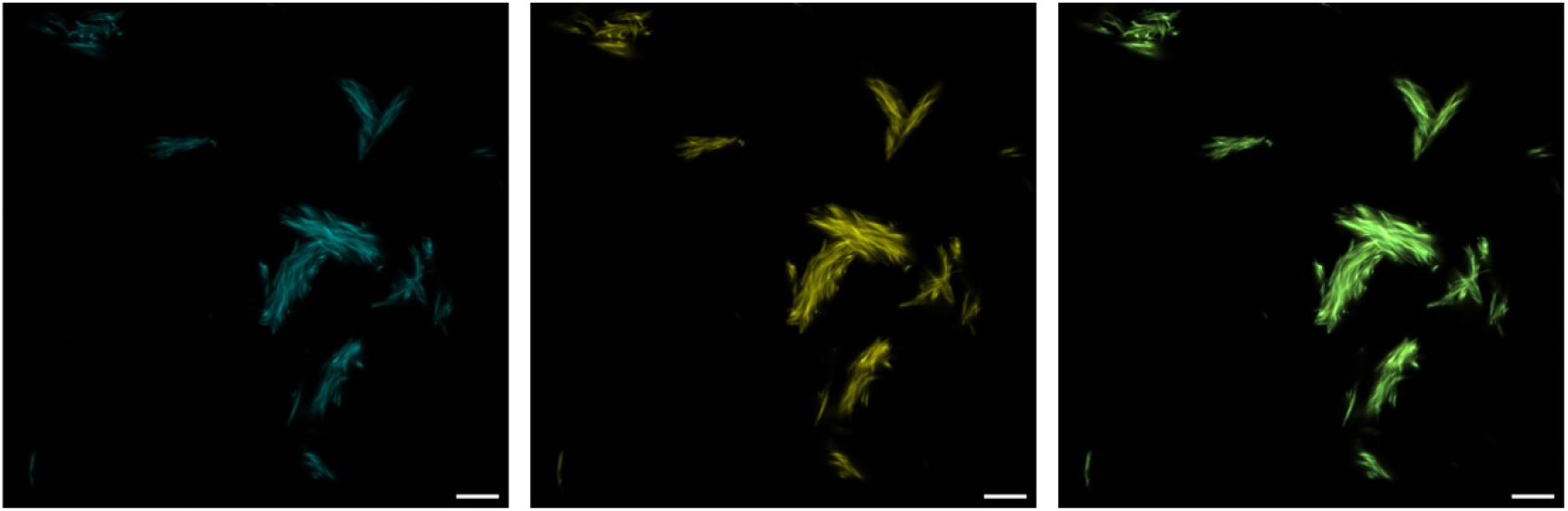
sfCyan and sfYellow colocalize to PAPf39 fibrils. 5 μM sfCyan and 5 μM sfYellow incubated with 30 μM preformed PAPf39 fibrils to confirm the simultaneous binding of monomeric fluorescent proteins. (A) sfCyan, (B) sfYellow, and (C) merged confocal microscopy channels. The scale bars are 5 μm.

**Figure S2:**
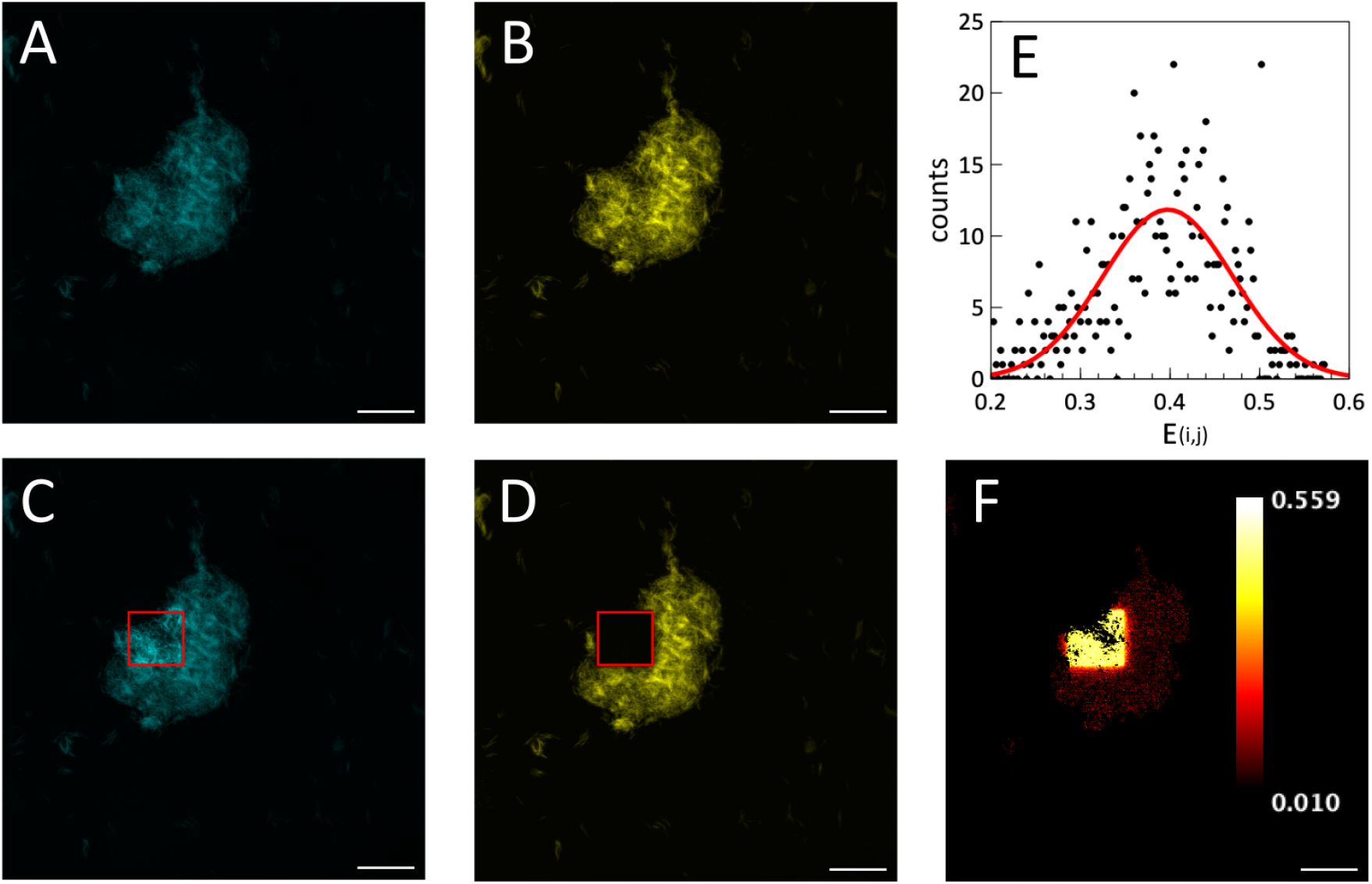
Example FRET acceptor photobleaching microscopy image set of PAPf39 fibrils incubated with YellowSC-STCyan. Images acquired using parameters to excite and detect only sfCyan (A and C; *λ*_ex_: 458 nm, *λ*_em_: 470-500 nm) or sfYellow (B and D; *λ*_ex_: 514 nm, *λ*_em_: 525-570 nm). Each channel was imaged prior to sfYellow bleaching (A and B) within a defined region of interest, and each channel was re-imaged after bleaching (C and D), with the sfYellow-depleted region shown in a red box. (E) Calculated FRET efficiency at each pixel used to generate a histogram of pixel counts per FRET efficiency value. The fit to equation 4 is shown as a red line. (F) Spatial representation of calculated FRET efficiency values at each pixel (value indicated by color given in calibration bar). Pixels outside the sfYellow-depleted region of interest are not included in further analysis. The scale bars are 25 μm.

**Figure S3:**
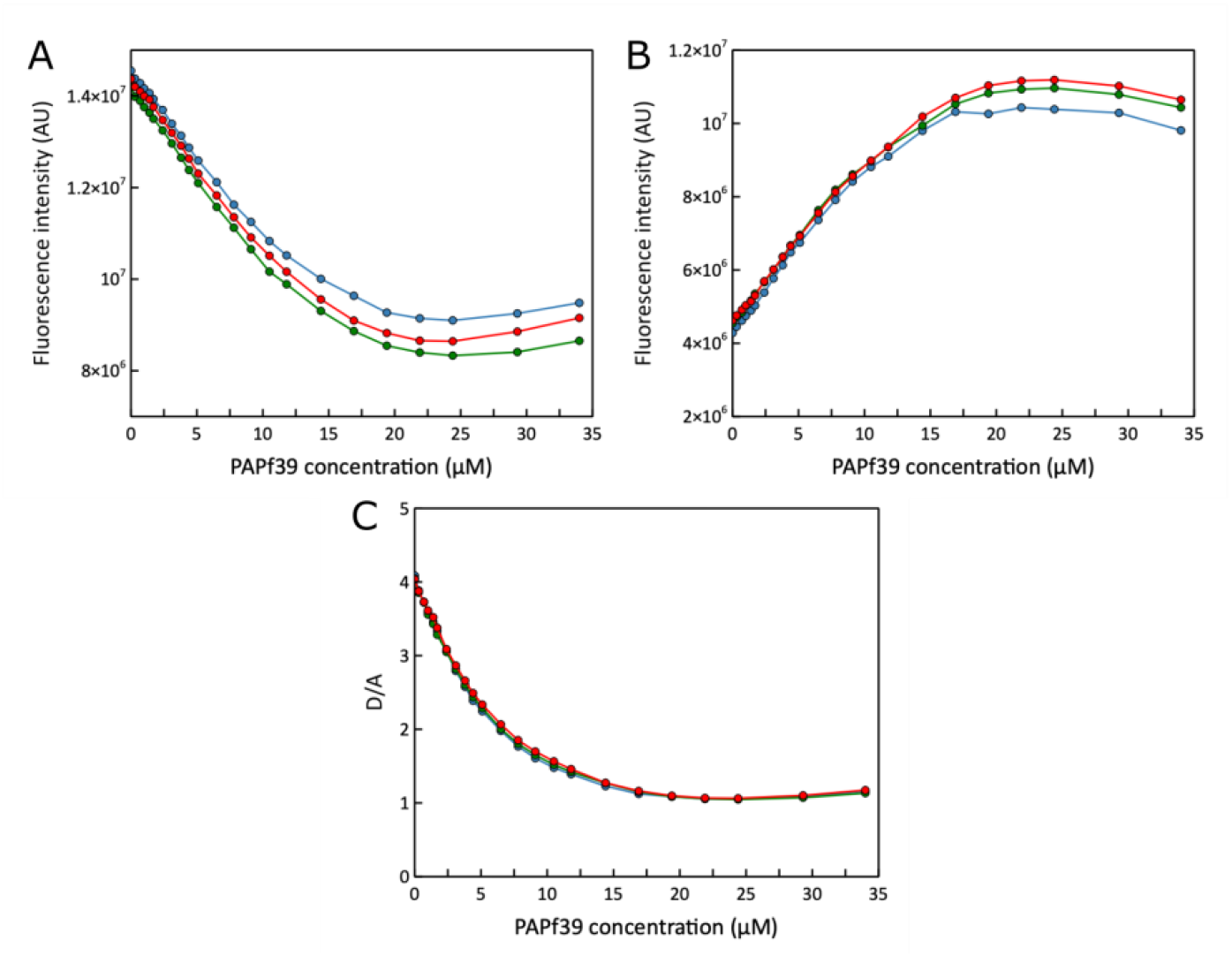
YellowSC-STCyan D/A metric provides more reproducible measurement than direct fluorescence intensities. Data from 3 independent replicate PAPf39 fibril titrations (red, green, and blue). (A) sfCyan and (B) sfYellow fluorescence intensities at their respective emission maxima (477 nm; 525 nm) derived from each unmixed YellowSC-STCyan spectrum as a function of PAPf39 fibril concentration. (C) D/A as a function of PAPf39 fibril concentration. Experimental variation due to small differences in fluorescent protein concentration and scattering at high fibril concentrations are masked in D/A calculation.

**Figure S4:**
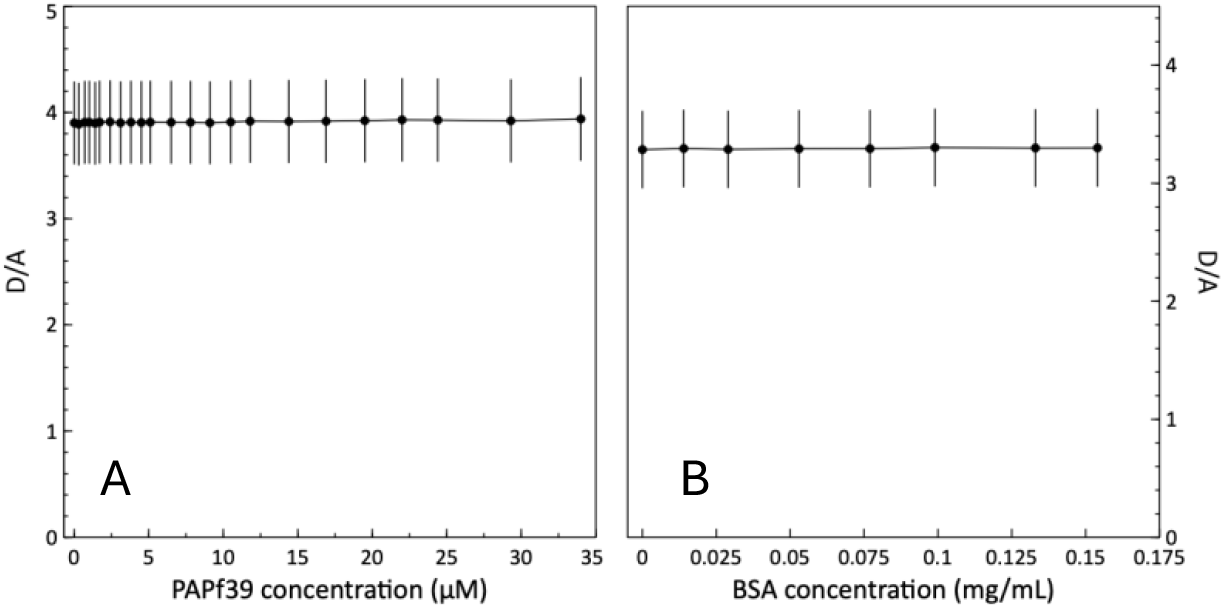
YellowSC-STCyan signal is insensitive to the presence of monomeric PAPf39 and amorphous protein aggregates. (A) YellowSC-STCyan D/A derived from raw emission spectra from 440-700 nm (*λ*_ex_ = 430 nm) as a function of monomeric PAPf39 peptide concentration over 2 replicates. (B) YellowSC-STCyan D/A derived from raw emission spectra from 440-700 nm (*λ*_ex_ = 430 nm) as a function of preformed BSA amorphous aggregate concentration over 3 replicates. Error bars on both panels represent 10% error.

**Figure S5:**
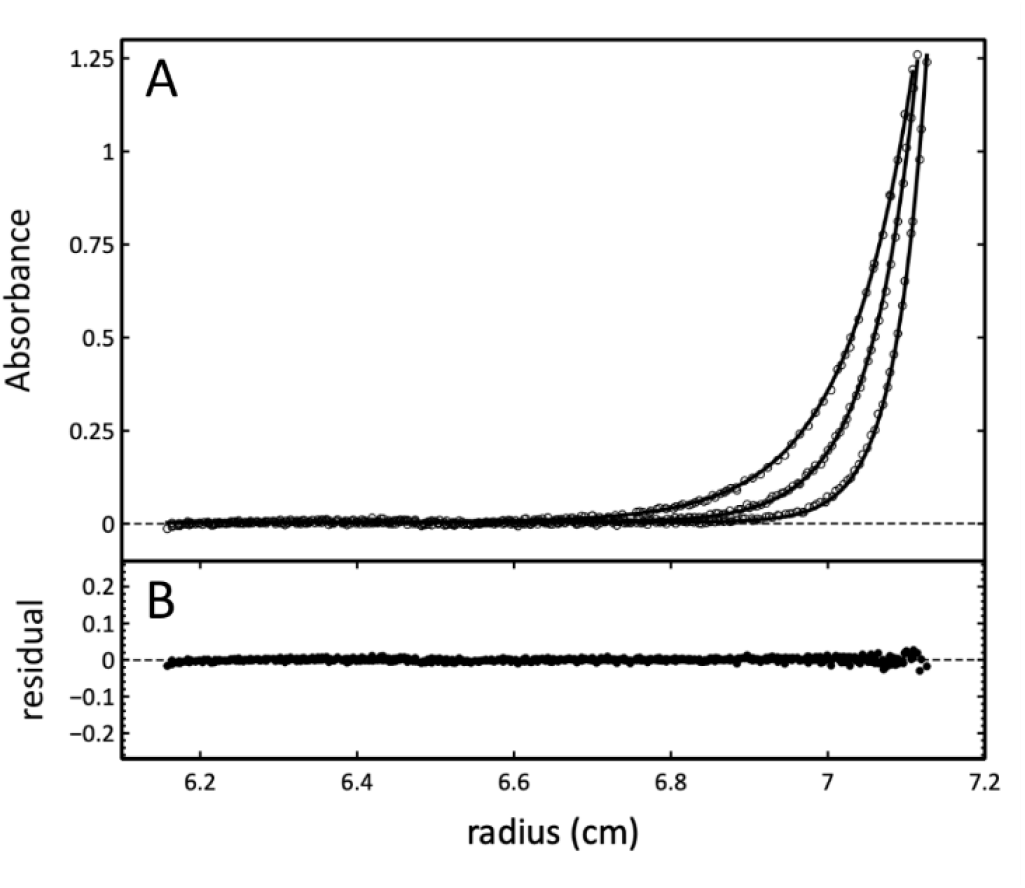
YellowSC-STCyan sedimentation equilibrium analytical ultracentrifugation data. (A) Raw data (open circles) and fits of each spectrum (solid lines) to Equation 1 from 15,000 rpm, 18,000 rpm, and 22,000 rpm scans. (B) Distribution of global residuals.

**Supplementary Table S1.**
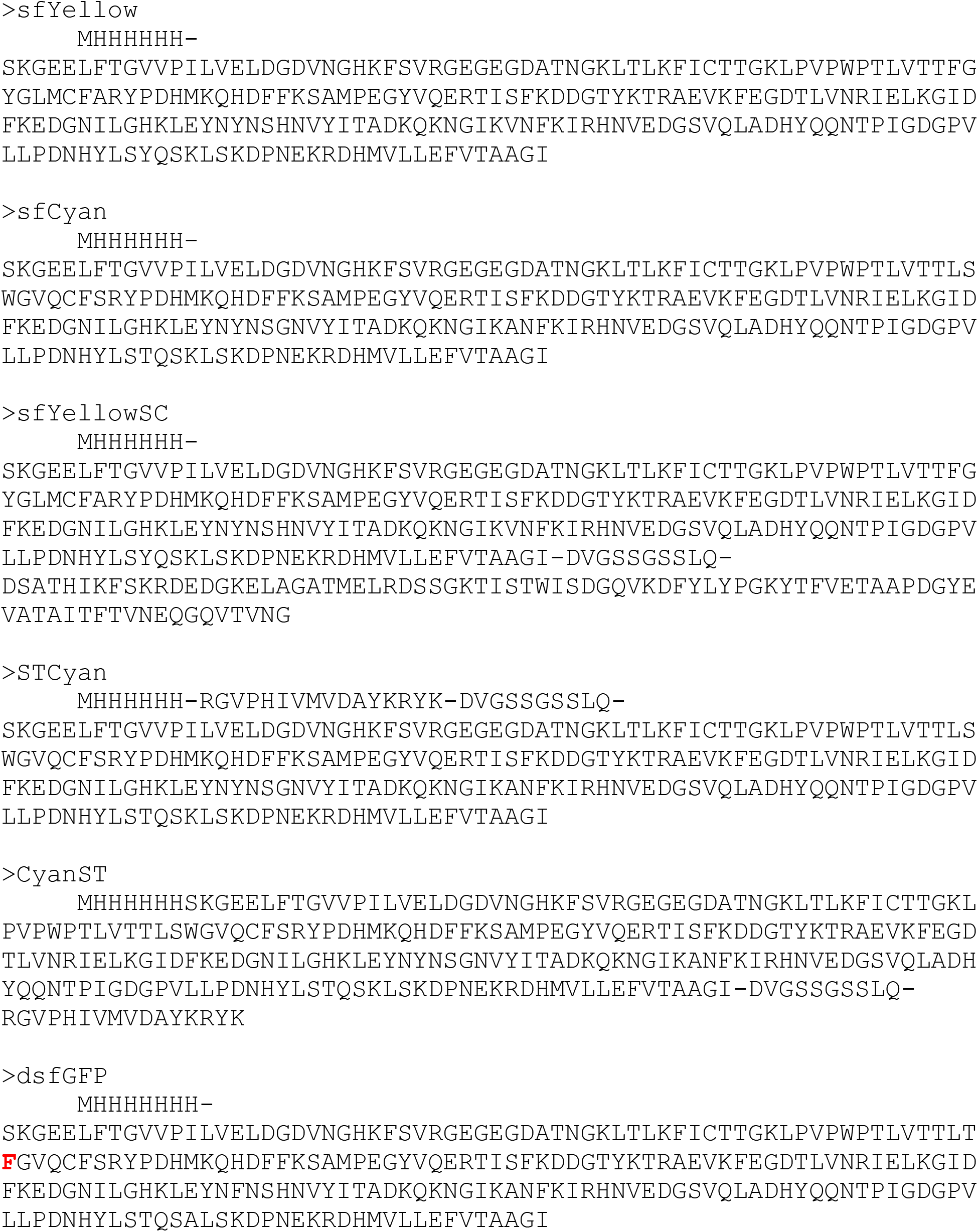
Protein sequences used in this work.

## Notes

### Competing Interest Statement

The authors have declared no competing interest.

